# Assessing species-specific neonicotinoid toxicity using cross-species chimeric nicotinic acetylcholine receptors in a *Drosophila* model

**DOI:** 10.1101/2025.03.19.644220

**Authors:** Anna Lassota, James J. L. Hodge, Matthias Soller

## Abstract

Nicotinic acetylcholine receptors (nAChRs) are ligand-gated ion channels and the main mediators of synaptic neurotransmission in the insect brain. In insects, nAChRs are pivotal for sensory processing, cognition and motor control, and are the primary target of neonicotinoid insecticides. Neonicotinoids are potent neurotoxins and pollinators such as honey bees are more sensitive and affected by extremely low sub-lethal doses. The pentameric nAChR channel is made up either of five α-subunits constituting five ligand-binding sites or a mixture of two to three α and β subunits constitute two to three ligand-binding sites. Of particular note, the honey bee nAChRα8 subunit is converted into a β subunit (nAChRβ2) in *Drosophila*, raising the question whether this α to β conversion makes flies less sensitive to neonicotinoids. To investigate species-specific aspects of neonicotinoid toxicity we CRISPR-Cas9 engineered a cross-species chimeric nAChR subunit by swapping the ligand-binding domain in *Drosophila* of nAChRβ2 with honey bee nAChRα8. Toxicity assessment by neonicotinoid thiamethoxam revealed significantly impaired motor functions in climbing and flight assays when comparing the α8/β2 chimeric channel to wild type or a β2 knock-out. However, both the α8/β2 chimeric channel and the β2 knock-out showed the same increased survival after neonicotinoid exposure compared to wild type flies. Combinatorial exposure to neonicotinoids also did not reveal differences. These findings highlight the critical role of nAChR subunit composition in motor control and demonstrate how subtle structural differences can profoundly impact motor function and pesticide response, offering new insights into the molecular mechanisms of neurotoxicity across species.

## Introduction

Pesticides are indispensable for protecting global agricultural yields by reducing crop losses caused by pests^1^. Among these, neonicotinoids are highly effective due to their broad-spectrum efficacy and systemic activity, which enable integration into plant tissues for targeted pest control, and versatility in application methods, such as seed coating, to reduce environmental contamination^2,3^. Neonicotinoids exhibit high toxicity to insects, but not to vertebrates, making them safer alternatives to other classes of chemical^4,5^. However, neonicotinoids do not distinguish between pests and beneficial insects, and harm beneficial insects like honey bees, which are critical pollinators for both wild flora and agricultural crops^6–8^. The widespread decline of pollinators, partially attributed to pesticide exposure, poses significant ecological and economic challenges, highlighting the need for pest-control strategies with reduced non-target effects.

Honey bees, wild bees and other pollinators are adversely affected by extremely low sub-lethal doses of neonicotinoids (2 ng/ml compared to the LC50 dose of 4.28 µg/ml) impairing foraging behaviour, cognition, navigation and weakening of the immune system, and reducing reproductive success and genetic diversity within a colony^6,9–20^. Neonicotinoids, which target cholinergic neurotransmission, inhibit mushroom body Kenyon cell activity^21^ and disrupt memory and olfactory sensory neuron activity in honey bees (*Apis mellifera*) and fruit flies (*Drosophila melanogaster*)^21–24^.

The sensitivity of insects to neonicotinoids varies considerably between species, with honey bees being highly susceptible and *Drosophila* more resistant. For example, the 24-hour lethal concentration 50% (LC50) of thiamethoxam (TMX) is 3.13 µg/ml for adult *Drosophila*^25^, whereas for honey bees it is 4.28 µg/ml^20^. This variation arises from differences in the repertoire of cytochrome P40 detoxification genes. *Drosophila* encodes 85 of these neonicotinoid-metabolising enzymes, whereas honey bees have only 46^26,27^. Additionally, honey bees have fewer antioxidant gene paralogs than *Drosophila*, which has an expanded antioxidant defence system. Since neonicotinoids can induce oxidative stress, this weaker antioxidant capacity may contribute to the greater sensitivity of bees compared to *Drosophila*^28^.

Another reason contributing to this species-specific susceptibility could be the composition of nicotinic acetylcholine receptors (nAChRs). nAChRs are essential for neurotransmission, synaptic plasticity, and neuronal development, making prominent targets for neonicotinoid pesticides^29–32^. These pentameric cys-loop ligand-gated cation channels consist of α and β subunits forming homo- or heteropentamers. Ligand binding pockets, located at α-α or α-β interfaces, consist of loops A, B, and C from an α subunit and loops D, E, and F from the adjacent α or β subunit^33–35^. Importantly, β subunits lack the ability to form functional homopentamers due to the absence of loops A, B, and C^33,35^. Consequently, the subunit composition of nAChRs receptor affect their properties and determines susceptibility to agonists, such as acetylcholine or neonicotinoids (Fig. 1a).

**Figure 1:**
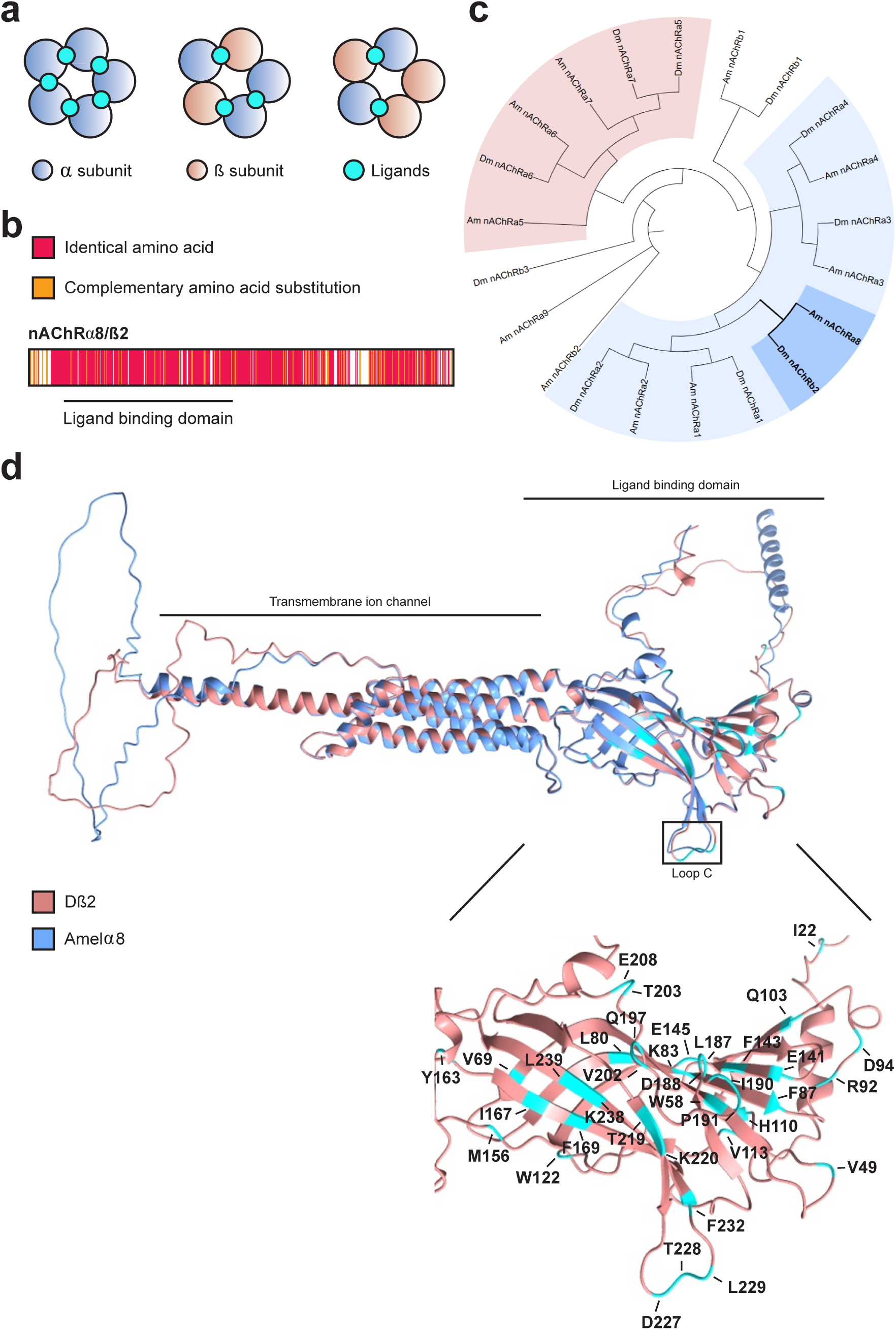
Highly conserved *Drosophila* β2 subunit originated from an alpha subunit in other insects. (a) Schematic representation of homo- and heteropentameric nAChR receptors in insects exhibiting different compositions of α (blue) and β (pink) subunits, with their active ligand binding sites represented by ligands (cyan). (b) Amino acid alignment of *D. melanogaster* β2 and honey bee Amelα8 subunits. Identical amino acids are marked as red, complementary amino acid substitutions are labelled orange, and non-complementary amino acid substitutions remained unmarked. The ligand binding domain is underlined. (c) Molecular phylogenetic analysis of all nAChR subunits present in *Drosophila* and honey bee with two major α subunits clades highlighted with light blue and pink (subunits highly conserved with mammalian α7). Species codes are Dm: *D. melanogaster* Amel: *A. mellifera*. (d) Structural alignment of *D. melanogaster* β2 (blue) and honey bee α8 (pink) with all amino acid differences in *D. melanogaster* ligand binding domain indicated (cyan).

Insects have relatively small nAChR gene families. *Drosophila melanogaster* encodes 10 subunits (Dα1-Dα7 and Dβ1-Dβ3), while the honey bee *Apis mellifera* encodes 11 (Amelα1-Amelα9 and Amelβ1-Amelβ2)^36^. α subunits are characterised by the presence of two adjacent cysteines in loop C of the ligand-binding domain (LBD), whereas β subunits lack these residues^35^. Notably, *Drosophila* encodes fewer α subunits compared to *Apis mellifera*. Instead, the *Drosophila* β2 subunit exhibits striking conservation with the honey bee Amelα8 subunit indicating a subunit conversion (Fig. 1b). However, the absence of loop C in *Drosophila* β2, which is essential for ion channel gating, highlights a structural divergence that may contribute to species-specific functional properties and differences in neonicotinoid susceptibility. Further, specific amino acid substitutions in nAChR subunits are known to impact insecticide susceptibility. For instance, *Drosophila* mutants lacking certain nAChR subunits or carrying the natural variant R81T in the β1 subunit from aphids exhibit a 10- to 100-fold increase in resistance to neonicotinoids^37–39^.

Here, we examine the evolutionary divergence and functional implications of nAChR subunits in *Drosophila melanogaster* and *Apis mellifera*. Phylogenetic analysis reveals that the *Drosophila* β2 subunit has evolved from an ancestral α subunit. To investigate whether this subunit conversion contributes to neonicotinoids sensitivity, we employed CRISPR-Cas9 gene-editing technology to generate chimeric *Drosophila* flies expressing a chimeric nAChR receptor. Specifically, we replaced the ligand-binding domain (LBD) of the *Drosophila* β2 subunit with that of the honey bee α8 subunit, providing a model to explore cross-species receptor functionality. Behavioural and survival assays revealed significant impairments in motor functions and altered sensitivity to insecticides, highlighting how structural differences in receptor subunits can influence pesticide response. Our results provide insights into the evolutionary trajectories of nAChR subunits and their roles in mediating neurotoxic compound interactions, establishing a valuable *Drosophila* model for investigating cross-species pesticide toxicity.

## Results

### Species-specific specialization of *Drosophila* β2 subunit during evolution

Insect *nAChRs* are classified into divergent subunits without clear orthologous relationships between species, alongside a subset of genes that are highly conserved across taxa^40^. Among these conserved subunits is the nAChRα8 subunit; however, it is absent in certain Dipteran species, including *D. melanogaster*. Instead, the *Drosophila* β2 subunit exhibits striking similarity to the honey bee α8 subunit, with 75% overall sequence identity and 92% identity within the LBD indicating that α8 converted in to the β2 subunit (Fig. 1b and Supplementary Fig. 1). Alphafold structural modelling further confirmed conservation of the protein secondary structure of these two subunits, particularly in the LBD, with the exception of the loop C (Fig. 1d)^41^.

Next, to investigate whether *Drosophila* β2 represents a divergent lineage from other β subunits or shares a closer evolutionary relationship with α subunits, we conducted a phylogenetic analysis of all nAChR subunits in *D. melanogaster* and *A. mellifera*. Strikingly, the analysis revealed that *Drosophila* β2 clusters within a clade predominantly composed of α subunits, including α1–α4 and α8, indicating that *Drosophila* β2 is more closely related to α subunits than to other β subunits (Fig. 1c). Interestingly, the honey bee β2 subunit exhibits significant divergence and does not closely group with any other subunits, suggesting a divergent evolutionary origin.

### *nAChR* subunits exhibit developmental dynamics and cell-specific distribution

To investigate the expression patterns of *nAChR* subunits, we analysed tissue specific expression data in adults, taking into account the enrichment values providing the gene abundance measure in a tissue relative to that in the whole fly^42^. The majority of *nAChR* subunits exhibits higher expression levels in females than in males (Fig. 2a). In both sexes, *nAChR* subunits are predominantly expressed in the head, brain and CNS, and thoraco-abdominal ganglion, with *Drosophila β2* being the most highly expressed subunit in the brain and CNS. In other tissues, however, the levels of expression differ between sexes, with *Drosophila β2* and *α7* being the most expressed subunits in the thoraco-abdominal ganglion of males and females, respectively. Furthermore, only two subunits, *Drosophila α4* and *β3*, are expressed across all tissues, with *Drosophila β3* showing notable expression in the heart. Specific differences were observed between sexes. For instance, in the male salivary glands, carcass, and crop, only *Drosophila α1*, *α2*, and *α3* were expressed, while *α4*, *β1*, and *β3* were restricted to females.

**Figure 2:**
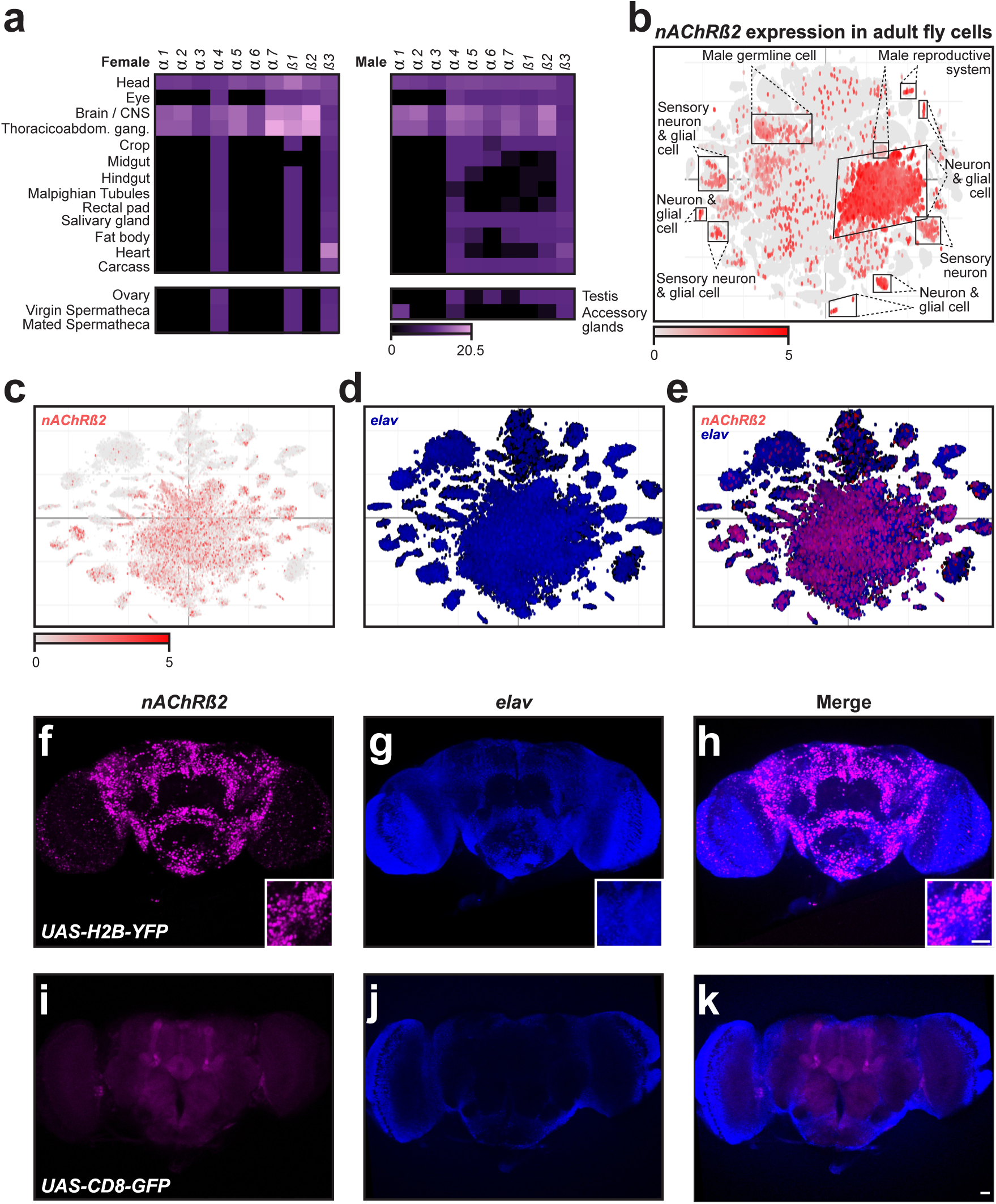
*Drosophila β2* is highly expressed in adult *Drosophila* brain and CNS but not in every cell. (a) Expression levels of *nAChR* subunits in female and male adult fly tissues. (b) Visualisation of single cell expression for the *nAChRβ2* gene indicated in red with annotation of the major cell clusters exhibiting *Dβ2* expression. (c-e) Visualisation of single cell expression for *nAChRβ2* (c) and neuronal marker *elav* (d) or both (e). (f-k) Expression of *Dβ2* (f, i), *elav* (g, j) and merged (h, k) in adult *Drosophila* brains with nuclear (f, h) and membrane-bound (i, k) *UAS* reporters. Scale bars are 20 µm.

Further, we investigated whether the expression of *nAChR* subunit changes during development. Larval tissue analysis revealed that *Drosophila α3* is not expressed in any tissue (Supplementary Fig. 2a), suggesting it is dispensable during development. In contrast, *Drosophila β3* was expressed in all larval tissues, with the garland cells showing the highest expression. The larval brain and CNS were the only tissues expressing all *nAChR* subunits, with *Drosophila β3* showing the highest levels. Overall, *nAChR* subunit expression is relatively low in most tissues in both larvae and adults (Fig. 2a and Supplementary Fig. 2a).

To refine the spatial expression of *β2* in *Drosophila* cells in more detail, we analysed expression in single cells^43^. In whole adult flies, *β2* was found to be expressed in various cell types, including neurons, sensory neurons, glial cells, as well as male germline and reproductive system, however, its expression is not ubiquitous (Fig. 2b). Notably, a prominent cluster of neurons classified as adult fly head CNS cells strongly expressed *β2*, suggesting it may have a significant role in neuronal circuits^23,44^. Since *β2* displayed the highest expression levels in the brain and CNS, we further analysed single cell expression data for the larval brain and whole adult fly head. In adults, *β2* expression was observed in distinct neuronal populations, contrasting with the broader expression of the neuronal marker *elav* (Fig. 2c-e)^45^. In contrast, larval *β2* was colocalised with *elav* in the majority of CNS neurons (Supplementary Fig. 1b-d)^46^.

To validate the expression patterns, we used *nAChRβ2^[2A-GAL4]^*allele which has a *T2A-GAL4* sequence fused at the 3’ end of the gene, resulting in GAL4 expression as a separate protein under the control of *nAChRβ2* regulation^47^. Therefore, we analysed both larval and adult brains using nuclear *UAS-Histone2B-YFP* or membrane-tethered *UAS-mCD8-GFP* reporters, co-staining neurons with anti-*elav* antibody. Consistent with our observations from the single cell expression analyses, *β2* expression was restricted to specific neuronal populations, confirming its non-ubiquitous distribution (Fig. 2f-k and Supplementary Fig. 1e-j). Furthermore, we noticed enhanced *β2* expression in Kenyon cells, particularly when a membrane-bound GFP reporter was used (Fig. 2i-k)^44^.

Together, these findings reveal that nAChR subunit expression is dynamically regulated during development and exhibits sex- and cell-specific specialisation, highlighting the functional diversity of these receptors in neural circuits and other tissues. Furthermore, the observed developmental shift in β2 expression, from widespread larval CNS expression to more specialised adult neuronal populations, reflects the transition from simpler larval neural circuits to the more specialised and complex adult CNS.

### Chimeric *Amelα8/Dβ2* ligand-binding domain flies are viable with impaired motor functions

Since the two adjacent cysteines in the loop C are essential for ligand binding in nAChRs, we hypothesised that this difference may influence neonicotinoid susceptibility, making honey bees more vulnerable. Therefore, we generated chimeric *nAChRβ2/α8* (*nAChRβ2^Amα8LBD^*) flies, where the *β2* LBD coding region in *Drosophila* was swapped with the *Amelα8* LBD coding sequence using CRISPR-Cas9 genome engineering (Fig. 3a and Supplementary Fig. 3). Successful recombinant flies were marked with *white+* selection marker, disrupting the LBD, therefore resulting in a null allele (*nAChRβ2^nullA^*). To restore the open reading frame (ORF) to generate a fly stock expressing the chimeric *nAChRβ2/α8* subunit (*nAChRβ2^Amα8LBD^*), the selection marker was then excised by PiggyBac transposase. Flies were validated by polymerase chain reaction (PCR), Sanger sequencing, and subsequently by whole genome sequencing.

**Figure 3:**
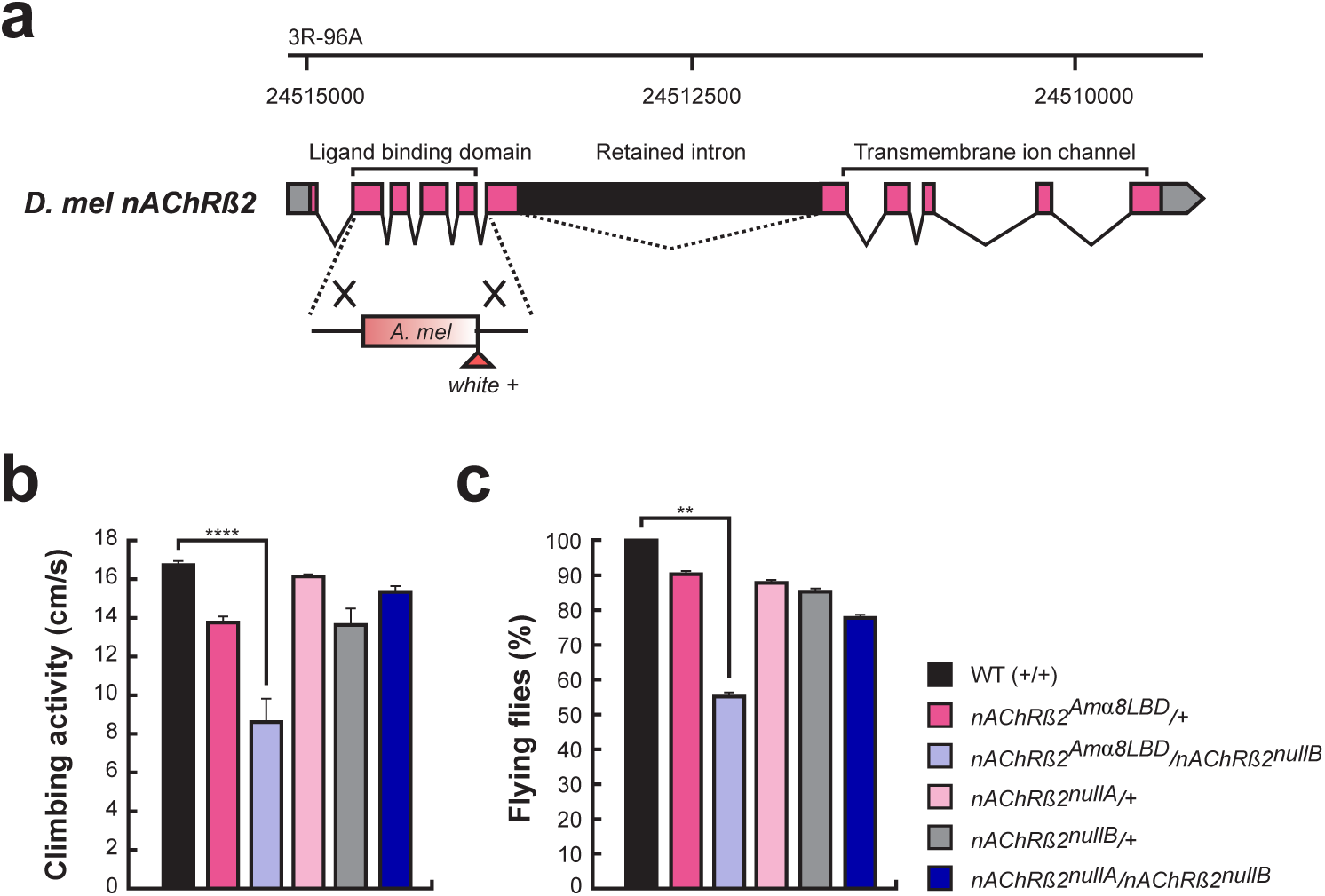
Chimeric *nAChRα8β2* flies exhibit locomotor defects. (a) Schematic *Dβ2* gene model with the ligand binding domain swap indicated with coding exons marked in pink. The *white+* selection marker was incorporated into the *nAChRβ2* gene along with the honey bee *nAChRα8* LBD encoding fragment, generating a *nAChRβ2^null^* allele. The reading frame can be restored by excising the PiggyBac transposon. (b) Climbing activity was assessed by negative geotaxis and is shown as means with the standard error from four biologically independent groups (n=20). Statistically significant differences are indicated by asterisks (\*\*\*\**p* ≤ 0.0001). (c) Flies of the indicated genotypes were tested for their flight ability shown as means with the standard error from four biologically independent groups (n=10). Statistically significant differences are indicated by asterisks (\*\**p* ≤ 0.01).

To normalise the genetic background, we used trans-heterozygotic flies for the *nAChRβ2^attP^* allele (*nAChRβ2^nullB^*) in which the coding region was replaced with an *attP* site, *3xP3-RFP* and a *loxP* site^47^, together with the *nAChRβ2^nullA^* allele. The chimeric *nAChRβ2^Amα8LBD^* mutants, as well as *nAChRβ2^nullA^* and *nAChRβ2^nullB^* flies are fully viable when crossed with a chromosomal deficiency (n = 453, 275, 421, respectively).

To assess motor abilities, we performed negative geotaxis assays (Fig. 3b) and evaluated flight ability (Fig 3c). In negative geotaxis assays the climbing ability of LBD swap mutant (*nAChRβ2^Amα8LBD^*/*nAChRβ2^nullB^*) was greatly reduced (*p* = 7.5*10^-7^) (Fig. 3b) compared to wild type (WT). We did not detect significant differences in the *β2* null mutant (*nAChRβ2^nullA^*/*nAChRβ2^nullB^*) nor the heterozygous mutant controls.

In flying ability assays, we observed the same, namely the LBD swap mutant (*nAChRβ2^Amα8LBD^*/*nAChRβ2^nullB^*) were impaired (*p* = 0.0066) (Fig. 3c) compared to control flies, but not the *β2* null mutant (*nAChRβ2^nullA^*/*nAChRβ2^nullB^*) or heterozygote mutant controls.

### Chimeric *Amelα8/Dβ2* ligand-binding domain flies display resistance to insecticides

To investigate the impact of substituting the *Drosophila* β2 LBD with the honey bee Amelα8 LBD on neonicotinoid susceptibility, we selected two well-characterised neonicotinoids, TMX and imidacloprid (IMI), both of which exhibit high toxicity to pollinators^5,39^. These compounds differ slightly in chemical structure and receptor specificity, with TMX being one of the most potent full agonists of *Drosophila* and honey bee α1, α2, β1 and β2 combinations when heterologously expressed in frog oocytes and assayed by patch-clamp. In contrast, IMI shows a stronger affinity for *Drosophila* β2-containing receptors compared to TMX^39,48^. Both compounds target multiple nAChR subunits, enhancing their versatility broad-spectrum toxicity^39^.

In addition, we tested flupyradifurone (FPF), a systemic butanolide compound that is chemically distinct from neonicotinoids and represents a novel alternative to these compounds in pest management^49^. While FPF is not primarily targeted at *Drosophila* β2 subunits^39^, its broad-spectrum activity and demonstrated toxicity to non-target pollinators, including bees^50,51^, make it an important candidate for assessing cross-resistance and potential off-target effects resulting from altered nAChR subunit composition.

First, we conducted survival analysis testing a range of TMX concentrations on WT flies (Supplementary Fig. 4a, b). Fly viability was measured over 6 days. Based on the survival curves, three different TMX concentrations were used for the toxicity experiments.

At 10 μM TMX, wild type flies had 43.75% survival after 144 h, while *nAChRβ2^Amα8LBD^/+*, *nAChRβ2^nullA^/+*, *nAChRβ2^nullB^/+*, *nAChRβ2^nullA^/nAChRβ2^nullB^*, and *nAChRβ2^Amα8LBD^*/*nAChRβ2^nullB^*flies remained viable (Fig. 4a). At 20 μM, survival rates declined progressively across all genotypes, with *nAChRβ2^nullA^*/*nAChRβ2^nullB^* mutants showing the highest survival rates, followed by *nAChRβ2^Amα8LBD^*/*nAChRβ2^nullB^* flies retaining more than 90% viability on day 6. This contrasted with *nAChRβ2^nullA^/+* (71.25%), *nAChRβ2^nullB^/+* (60%), *nAChRβ2^Amα8LBD^/+* (35%), and WT (11.25%) (Fig. 4b). At 40 μM, similar trend was observed, although the viability of all flies was decreased below 50%. *nAChRβ2^nullA^*/*nAChRβ2^nullB^*mutants retained by 48.75% day 6, followed by *nAChRβ2^Amα8LBD^*/*nAChRβ2^nullB^*(41.25%), *nAChRβ2^nullA^/+* (35%), *nAChRβ2^nullB^/+* (16.25%), *nAChRβ2^Amα8LBD^/+* (3.33%), while WT flies showed 0% survival by day 4 (Fig. 4c). Interestingly, apart from the trans-heterozygous *β^nullA^*/*β^nullB^* flies, all genotypes experienced a sharp drop in viability between the 1^st^ and 2^nd^ day of treatment, particularly *nAChRβ2^Amα8LBD^/+* dropping from 62.5% to 30% and WT failing from 79% to 20%.

**Figure 4:**
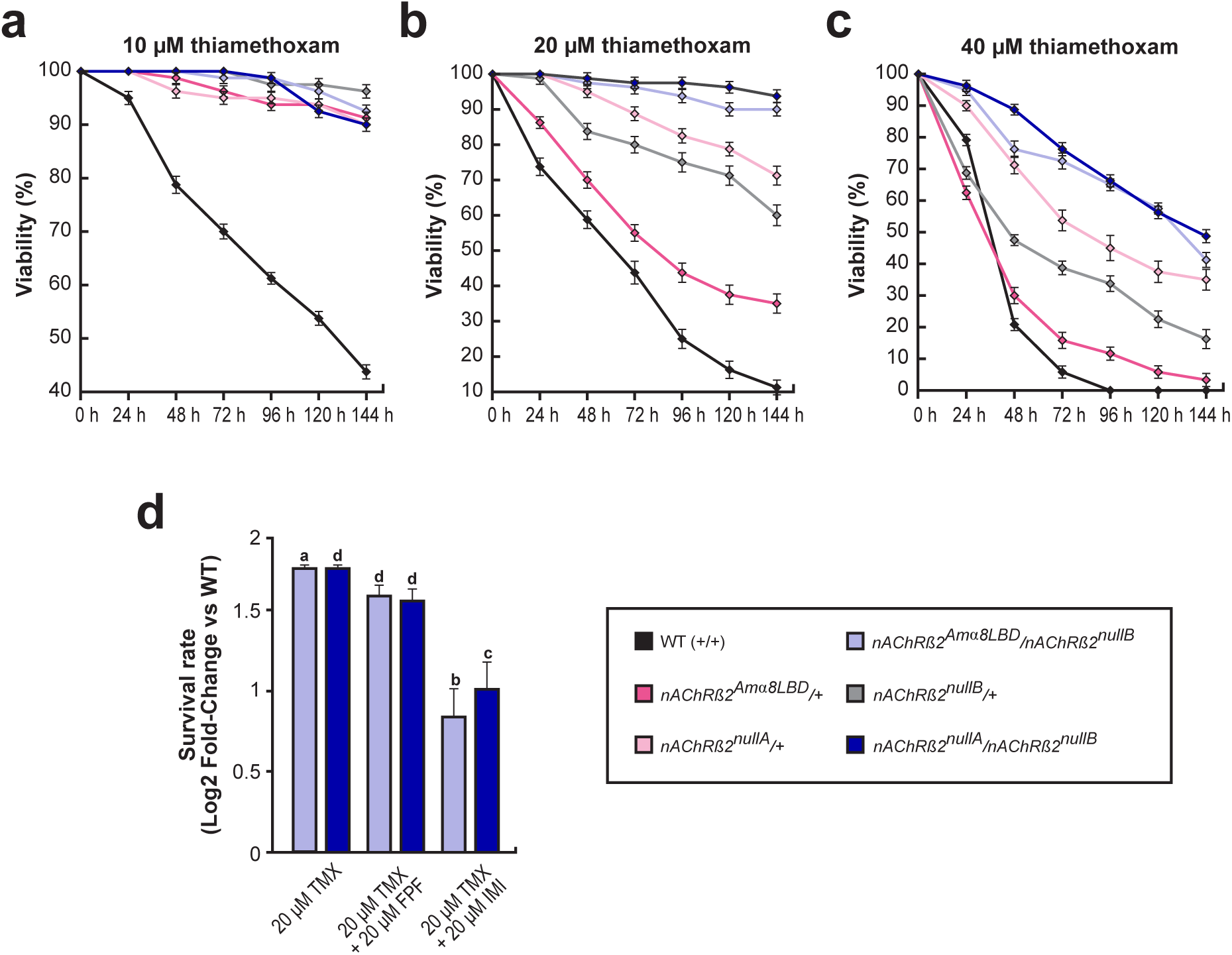
The α8 ligand binding domain does not sensitize flies to neonicotinoids. (a-c) Flies viability exposed to 10 µM (a), 20 µM (b) and 40 µM (c) thiamethoxam measured every 24 h for 6 days is shown as mean with standard error from four biologically independent groups (n = 20). (d) Survival rate of *nAChRβ2^Amα8LBD^*/*nAChRβ2^nullA^*and *nAChRβ2^nullA^*/*nAChRβ2^nullB^* mutants compared to WT flies subjected to different insecticides combinations. The log2 fold-change shown as mean with standard error from four biologically independent groups (n = 20) represents the viability of flies after six days of treatment with thiamethoxam (TMX) or TMX mixed with imidacloprid (IMI) or flupyradifurone (FPF) in equimolar ratios. Statistically significant differences are indicated with letters (a = \**p* ≤ 0.5, b = \*\**p* ≤ 0.01, c = \*\*\**p* ≤ 0.001, d = \*\*\*\**p* ≤ 0.0001).

While both neonicotinoids and butenolides act as nAChR agonists, bind to the same site and cause prolonged activation, they vary in the binding dynamics, receptor subtype preferences, and potential to develop resistance^52^. To assess whether combining neonicotinoids and butanolides results in additive or synergistic toxicity, we subjected flies to a mixture of TMX (20 μM) with either FPF (20 μM) or IMI (20 μM). In nature, pollinators and insects are frequently exposed to multiple insecticides rather than a single compound, making it crucial to understand potential interactions and cross-resistance effects^53,54^. The TMX concentration was chosen based on prior survival analyses, where double mutant flies retained > 90% viability (Fig. 4b), while WT flies showed significant susceptibility. We tested chimeric *nAChRβ2^Amα8LBD^*/*nAChRβ2^nullB^*flies and *nAChRβ2^nullA^*/*nAChRβ2^nullB^* mutants to determine whether altered receptor composition influences responses to dual insecticide exposure. Flies were treated for six days, and viability on day six was analysed as a log-fold change relative to WT control and TMX-only treatment (Fig. 4d). Notably, exposure to TMX in combination with either FPF or IMI resulted in a slight increase in survival rates. However, the combined treatment did not significantly alter overall insecticide susceptibility. As expected, *nAChRβ2^Amα8LBD^*/*nAChRβ2^nullB^*flies and *nAChRβ2^nullA^*/*nAChRβ2^nullB^* mutants flies exhibited significantly higher resistance compared to WT across all conditions.

Taken together, our findings demonstrate that *nAChRβ2^nullA^*/*nAChRβ2^nullB^* null mutants as well as the LBD swap mutants *nAChRβ2^Amα8LBD^/nAChRβ2^nullB^* flies exhibit higher survival rates upon exposure to TMX, reinforcing the role of *Drosophila* β2 in neonicotinoid susceptibility. Further, exposure to combined neonicotinoid and butanolide treatments did not significantly alter overall lethality compared to TMX treatment on its own, indicating that sub-lethal doses of these nAChR-targeting insecticides do not act synergistically in this context. The observed resistance of chimeric and β2^null^ mutants further highlights the functional divergence of Amelα8 in modifying nAChR-mediated insecticide susceptibility.

## Discussion

The agricultural industry relies on pesticides to protect crops and maintain a stable food supply for a growing population. Among these, neonicotinoids are the most widely used insecticides due to their high efficacy and systemic action, reducing the need for repeated applications. However, residues in nectar and pollen pose a significant risk to pollinators. Despite their well-documented toxicity to bees and other beneficial insects, neonicotinoids remain the dominant insecticides worldwide^55^, highlighting the urgent need to mitigate their ecological impact.

Beyond their effects on pollinators, neonicotinoids exhibit cross-species variability in toxicity, with *Drosophila melanogaster* showing notably higher resistance than honey bees, despite both species having homologous nAChRs^48^. The mechanisms underlying this differential sensitivity remain unclear but are likely influenced by differences in receptor composition, metabolism, or detoxification pathways^26,28^. To investigate species-specific differences in neonicotinoid sensitivity, we established a *Drosophila* model where we swapped the LBD of *D. melanogaster* β2 with honey bee Amelα8 subunit. The LBD swap mutant did not reveal increased sensitivity to neonicotinoids compared to the *Drosophila* β2 null mutant. However, the LBD swap mutant displayed motor function deficits indicating that this chimeric receptor is functional. Minor structural changes in receptor subunits can profoundly impact pharmacological properties^33,35,38,39^ and could account for the altered neonicotinoid responses observed in LBD chimeric mutant flies.

Previous studies in honey bees revealed tissue-specific expression of nAChR subunits, including α2, α8 and β1 in Kenyon cells, while α7 was restricted to antennal lobe neurons^56^. A similar functional diversity is observed in *Drosophila*, where nAChR subunits like α1 and α6 contribute to dendrite morphogenesis and synaptic transmission within larval visual circuits^57^. The absence of α3 expression in *Drosophila* larvae, combined with reduced nAChR subunit expression during early developmental stages, suggests that receptor composition undergoes developmental reprogramming to support the formation of complex adult neural circuits. This dynamic tissue-, sex-, and stage-specific expression underscores the diverse roles of nAChRs, including potential adaptations for behaviours such as reproduction^58,59^. Interestingly, in the cockroach *Periplaneta americana*, subchronic exposure to sublethal doses of imidacloprid induces a significant decrease in α2 mRNA expression, which correlates with a reduced sensitivity to the insecticide^60^, highlighting potential adaptive changes in nAChR subunit composition even in fully developed insects.

Furthermore, high *Drosophila* β2 expression levels in CNS-specific neuronal populations, including Kenyon cells, suggest its involvement in higher-order neuronal functions such as learning and memory^44^. Similarly, honey bee α8 is enriched in honey bee mushroom bodies^36,61,62^, suggesting a conserved role for *Drosophila* β2 and honey bee α8 in regulating mushroom body function across insect taxa, despite species-specific specialisations. Notably, it has been shown that exposure to neonicotinoid insecticides affects cell activity in mushroom body, and impairs learning and memory in insects, including *Drosophila* and honey bees^21–23,63^, further highlighting the critical role of nAChRs in cognitive functions and suggesting that disruption of these receptors can have detrimental effects on neuronal processes related to behaviour and learning.

Pharmacological and genetic studies have identified α1, α2, β1, and β2 as the primary mediators of neonicotinoid toxicity^40,48,64^. Here, we generated a cross-species chimeric nAChR using CRISPR-Cas9 genome engineering to evaluate neonicotinoid sensitivity in a *Drosophila* model. Our study reveals that the *Drosophila* β2 conversion of honey bee α8 is not a main determine of neonicotinoid sensitivity requiring to expand this approach to the other main mediators of neonicotinoid toxicity. Our study establishes a framework for investigating molecular determinants of neonicotinoid sensitivity of insects and vertebrates nAChR using a cross-species chimeric nAChR *Drosophila* model. This model can contribute to the design of next generation of insecticides for enhanced species-specific responses with implications for pollinator conservation and pesticide safety for vertebrates including humans.

## Materials and methods

### Fly stocks, genetics, immunostaining of tissues and imaging

*D. melanogaster* CantonS and w^1118^ were used as the wild type control. *nAChRß2^attP^* (BDSC 84545) and *nAChRβ2^[2A-GAL4]^* (BDSC 84666) stocks were described previously ^47^ and were together with the chromosomal deficiency (BDSC 24996) from Bloomington. Fly crosses were maintained at 25°C in plastic vials containing 10 ml of a standard cornmeal/yeast-rich medium (1% agar, 2% yeast, 7% dextrose, 8% cornmeal w/v and 2% Nipagin from a 10% solution in ethanol) with a 12:12 h light–dark cycle.

Third instar wandering larvae and adult brains from the progeny of *UAS-Histone2B-YFP/+*;*nAChRβ2^[2A-GAL4]^/+* and *UAS-mCD8-GFP/+*;*nAChRβ2^[2A-GAL4]^/+* were dissected in phosphate buffered saline (PBS) and fixed in 4% paraformaldehyde in PBT (PBS with 0.1% TritonTM X-100 (Sigma-Aldrich, T8787)) for 30 min, followed by washes in PBT 3 × 15 min. Samples were incubated overnight at 4°C with primary mouse anti-*elav* antibodies (MAb 7D, 1:20)^65^, followed by secondary antibodies conjugated with Alexa Fluor 546 again overnight at 4°C. Samples were counterstained with DAPI (1:1000), mounted in Vectashield (Vector Labs), scanned with Leica SP8, and processed using FIJI.

### Sequence analysis and single cell expression data visualisation

Amino acid sequences were aligned using ClustalW with Megalin (DNAstar) or with MAFFT^66^ and phylogenetic trees were generated in NGPhylogeny.fr^67–69^ and visualised using iTOL^70^.

Raw whole-genome sequencing reads were assessed for quality using FastQC, followed by adapter trimming and quality filtering with Trim Galore. High-quality reads were aligned to the *Drosophila melanogaster* (dm6) and *Apis mellifera* (Amel_HAv3.1) reference genomes using BBMap. Mapping quality and coverage were evaluated, and alignments were visualized using IGV (Integrative Genomics Viewer).

Single cell expression data was visualised as t-distributed stochastic neighbour embedding (tSNE) from the 10x Stringent dataset in Scope and ASAP^71^.

### Generation of chimeric *nAChRα8β2* flies

Two single guide RNAs (sgRNAs) flanking the ligand binding domain *Dβ2* were designed using PlatinumCRISPr^72^ (Supplementary Fig. 3a, b) and cloned into *pUC-3GLA* using the following primers: nACHr8 sgRNA left F1 (AAGATATCCGGGTGAACTTCGCTTATTGGAGCTAGGAAAGGTTTTAGAGCTAGA AATAGC) and nACHr8 sgRNA right R1 (GCTATTTCTAGCTCTAAAACCACATGGCACAATCAAATTCGACGTTAAATTGAAA ATAGG) as described previously^72^. Chimeric nAChR subunits were constructed by combining sequences of *nAChRα8* from *A. mellifera* and *nAChRβ2* from *D. melanogaster* in three steps (Supplementary Fig. 3c-e). Fragments used for generating the chimeric nAChR subunit were synthesised using PCR with primers Bee alpha8 F2 (GGGAGTGGTCATTGCCATCTCAACCCTATATAAATTTG) and Bee alpha8 R2 (GGAATGTAATACCCACGCAAGGAATTATTAAG), followed by Bee alpha8 F1 (GCCAGTGAATTCGAGCTCGGTACCGCAGCTAGCGAAGCAAATCCTGACACAAAGAGACTTTATGATGAC) and Bee alpha8 R1 (TTTACGCAGACTATCTTTCTAGGGTTAACCGTATAGAATAATGTTTTTCTTCGCATTG) for Amelα8 ligand binding domain, Dmel nAChRb left F1 (CGGCCAGTGAATTCGAGCTCGGTACCGATCCTTTTAGATAAAACATTTAGGAGCT ATC) and nAChRsgmut left R1 (CTCTTTGTGTCAGGATTTGCTTCGAAACTCACTGGAGCCGTGAGAGGAGGGGTAATTCAGGGGAAAAACAGAGAAAAATGCC) for Dβ2 left homology arm, and nAChRsgmut right F1 (ATTTTACGCATGATTATCTTTAACGTACGTCACAATATGATTATCTTTCTAGGGTTAATCTGATTGTGCCCTGCGTAGCTTTAACATTCC) and Dmel alpha8 right R1 (GCATGCCTGCAGGTCGACTCTAGAGGATCCTGATAGTTGCTGCTGCGAATGCGG AGCTG) for Dβ2 right homology arm, ensuring sgRNA sites in both homology arms are mutated. Mutations were introduced based on conservation of the region between closely related *Drosophila* species analysed using the UCSC genome browser (https://genome.ucsc.edu) as described previously^73^. Fragments were then cloned stepwise into the *pUC19 pBac w+* [accession number: PV267745] containing *white +* marker, disrupting the LBD, therefore resulting in *nAChRβ2^null^* flies. The selection marker was flanked with inverted terminal repeats inserted at a TTAA PiggyBac motifs for the later scarless excision by PiggyBac transposase.

The *pUC19 pBac w+ Amelα8/Dβ2* plasmid was treated with the nicking endonuclease *Nb.Bts*I (NEB, R0707S) prior injection. Transgenic lines were generated by injection into a GFP nosCas9 carrying flies (kindly provided by FlyORF). Positive transformants were identified by red eye colour.

For *Drosophila* whole genome sequencing, large fragment DNA was extracted with the Quick-DNA Tissue/Insect Miniprep Kit (Zymogene) according to the manufacturers’ instructions but replacing the BashingBead lysis step with cracking flies in liquid nitrogen followed by gentle homogenization with a pestle in the provided BashingBead Buffer. Illumina sequencing was done by Novogene and the sequence submitted to GEO [PRJNA1233772].

### Locomotive assessment

Negative geotaxis experiments were done as previously described^74^. Briefly, two to five day-old flies of both sexes were collected with CO_2_ anaesthesia and grouped in sets of 20. Flies were allowed to recover for a day and then placed in two inverted fly vials (19 cm). Flies were tapped to the bottom, and their climbing behaviour was recorded on video. Every 5 s for a total of 30 s, the distance climbed by each fly was measured and recorded. The data was collected for four biological replicates.

For flying ability assessment, two to five day-old flies of both sexes were collected with CO_2_ anaesthesia and grouped in sets of 10. Flies were allowed to recover for a day before the flying test. Then, flies were tapped to a flat surface, and their ability to fly was recorded on video. Flies were observed over a 30 s period. At the end of 30 s, the number of flies that successfully flew away was recorded. The data was collected for four biological replicates.

### Insecticide toxicity assay

Two to five day-old flies of both sexes, grouped in sets of 20, were exposed to various concentrations of thiamethoxam (Sigma-Aldrich, 37924), or a combination of 20 µM of thiamethoxam with 20 µM imidacloprid (Thermo Fisher, 466752500) or 20 µM flupyradifurone (MCE, HY-145295). The insecticides were dissolved in 0.01% acetone and added on top of the standard fly food, and flies were added to the vials after 24 h. Exposure concentrations ranged from 1.25 µM to 80 µM for survival curve analysis (Supplementary Fig. 4) and from 5 µM to 40 µM for testing all genotypes. Flies were exposed for 6 days, and survival was assessed every 24 h.

### Statistical analysis

Statistical analyses were carried out using GraphPad Prism 9, applying one-way ANOVA for behavioural assays or two-way ANOVA for the toxicity assays followed by Tukey’s test for pairwise comparisons. The data represent a minimum of three independent replicates conducted on separate days.

## Acknowledgments

We thank the Bloomington stock centre and FlyORF for fly lines, FlyORF for *Drosophila* injection service for embryo injections, and Al Handler for the PiggyBac plasmid.

## Authors’ contributions

A.L. performed all experiments and analysed data. M.S. and J.J.L.H. conceptualised the project. A.L. wrote the manuscript and M.S. and J.J.L.H. reviewed it. All authors read and approved the final manuscript.

## Availability of data and materials

The data generated or analysed during this study are included in the supplementary information in the supplementary information files. Whole genome sequencing data have been deposited at GEO under the accession number PRJNA1233772. The sequence for the *pUC19 pBac w+* has been deposited in GenBank under the accession number PV267745 and the plasmid is available from Addgene and the European plasmid repository.

## Declarations

### Ethics approval and consent to participate

Not applicable.

### Consent for publication

Not applicable.

