## Supplementary Materials for "Assessing species-specific neonicotinoid toxicity using cross-species chimeric nicotinic acetylcholine receptors in a *Drosophila* model"

#### for

**Supplementary Figure 1: Amino acid alignment of *Drosophila* Dβ2 and honey bee Amelα8**

Amino acid alignment of *D. melanogaster* Dβ2 and honey bee Amelα8 subunits. Identical amino acids are marked as red, complementary amino acid substitutions are labelled orange accordingly to the BLOSUM-62 substitution matrix which is a quantitative approach for assessing whether an amino acid substitution is conservative or nonconservative. The non-complementary amino acid substitutions remained unmarked. The ligand binding domain is underlined, and the loop C is marked with a black box.

**Supplementary Figure 2: *Dβ2* is highly expressed in larval *Drosophila* brain and CNS but not in every cell**

(a) Expression of *nAChR* subunits in larval tissues.

(b-d) Uniform manifold approximation and projection for dimension reduction (UMAP) projections of the *Dβ2* (b), *elav* (c) or both (d) expression in the larval brain and CNS cells based on Avalos *et al.* single cell datasets.

(e-j) Expression of *Dβ2* (e, h), *elav* (f, i) and merged (g, j) in larval *Drosophila* brains with nuclear (e, g) and membrane-bound (h, j) *UAS* reporters. Scale bar is 20 μm.

**Supplementary Figure 3: Chimeric *nAChRβ2/α8* construct cloning strategy and sgRNA design**

(a, b) Sequences and predicted secondary structures of single guide RNAs targeting the homology arm upstream the ligand binding domain (LBD) termed "left" (a) and downstream the LBD termed "right" (b). PAM sequence and additional G for enhanced base-pairing are highlighted in red.

(c, d) The sgRNAs sequences were mutated in the construct as indicated by red nucleotides and amino acid, as well as grey nucleotides in the sequence upstream of the left sgRNA. The lower- and upper-case nucleotides represent intron and exons, respectively. I22V mutation was introduced to mimic the amino acid residue in the honey bee LBD sequence (c).

(e) Schematic representation of step-by-step construct generation. The restriction sites used for cloning the next fragment into the vector are indicated. The final construct contains honey bee LBD, *D. melanogaster* left and right homology arms (LHA and RHA) with mutated sgRNAs, as well as the *white*+ (*w*+) selection marker flanked with inverted terminal repeats and TTAA sequences for its later scarless excision by PiggyBac transposase to restore the reading frame of the nAChR receptor.

**Supplementary Figure 4: Effect of various insecticides concentrations on *Drosophila* viability.**

(a) Survival curves of WT flies after 6 days of treatment with increasing thiamethoxam concentrations. Viable flies were counted every 24 h.

(d) Viability of WT flies exposed to increasing thiamethoxam concentrations after 24 h. The dotted line denotes when the survival falls below 50% (LC50).

Data are presented as mean ± SD from three biologically independent groups (n = 20). Error bars represent standard errors.

### Supplementary Figure 1

**a** **nAChRα8/ß2**

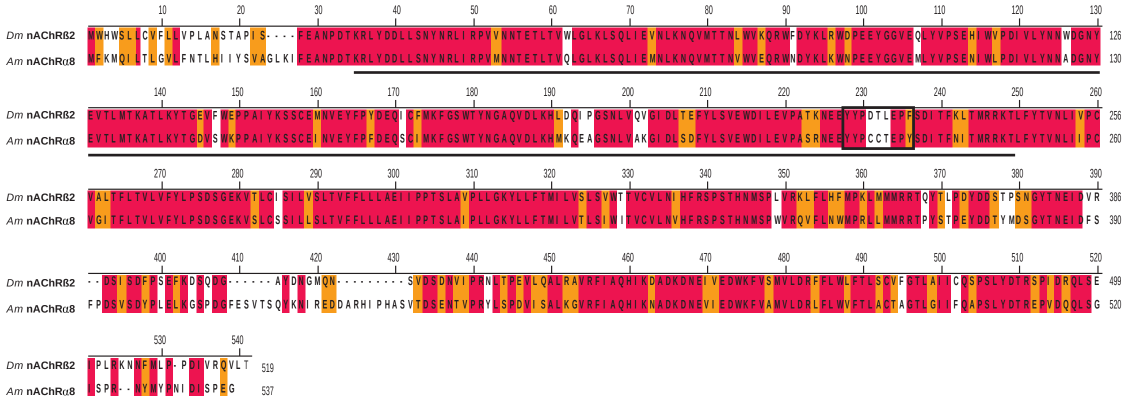

### Supplementary Figure 2

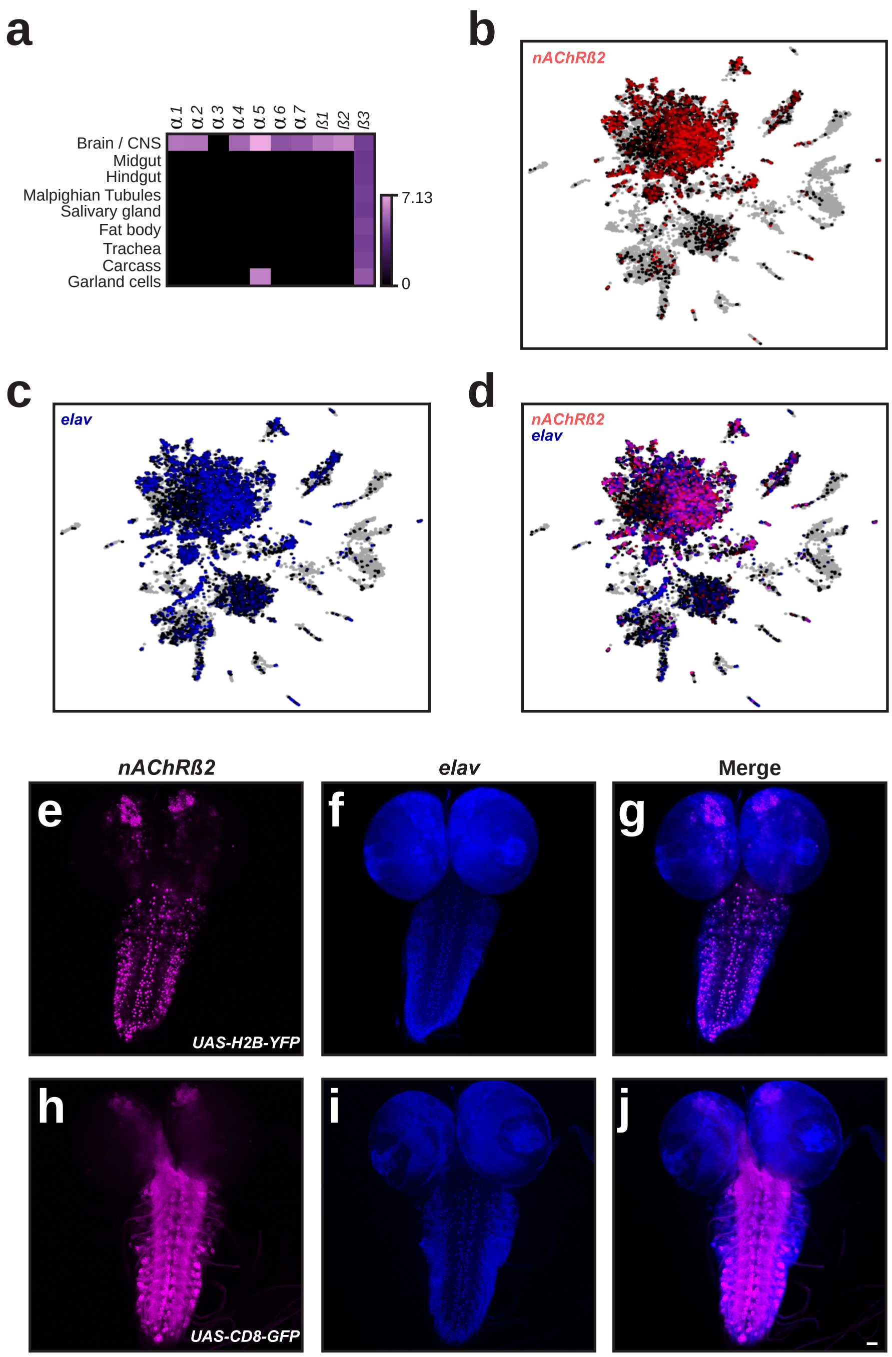

### Supplementary Figure 3

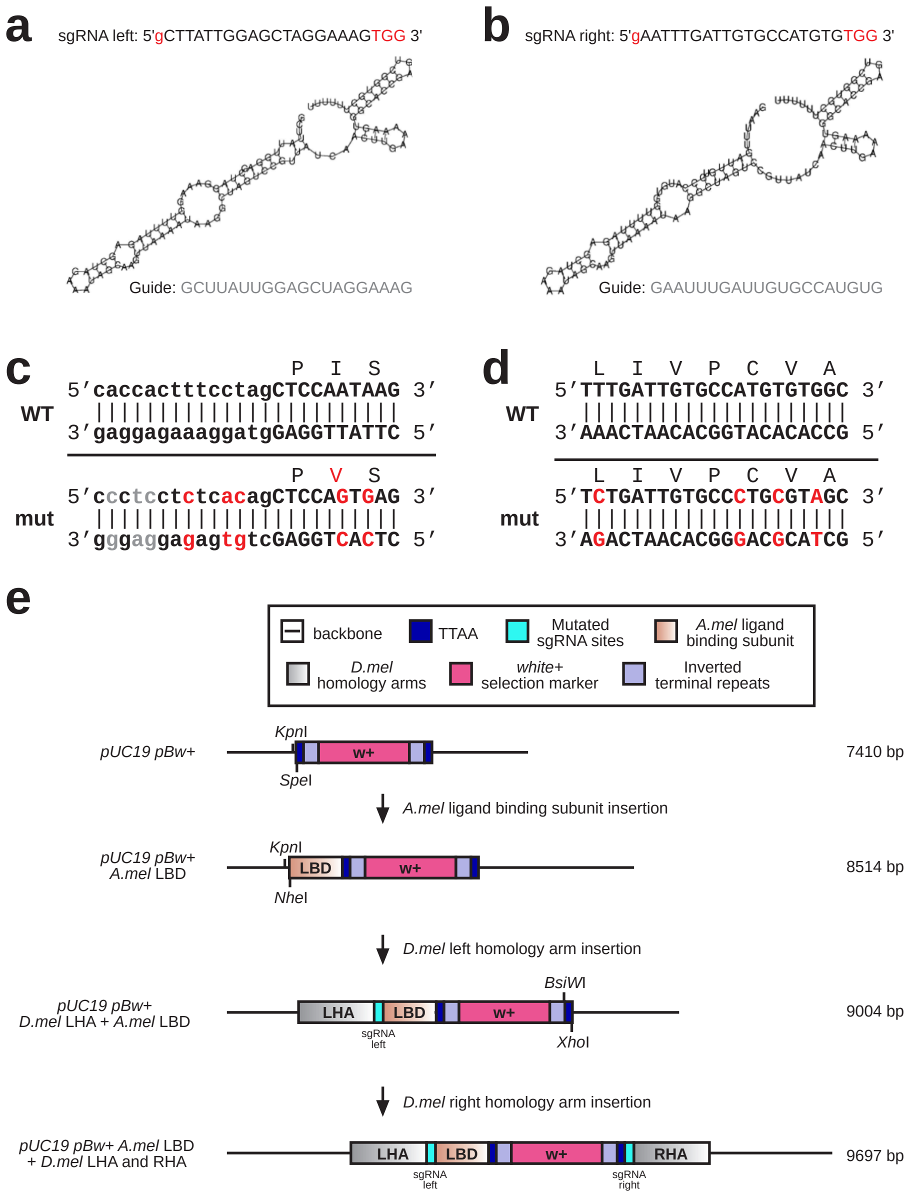

### Supplementary Figure 4

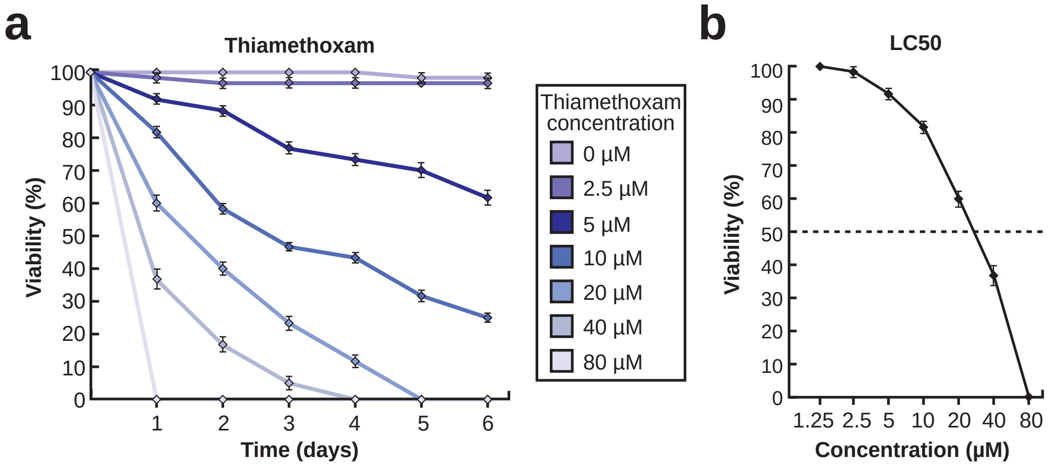
